# Mitochondrial DNA has strong selective effects across the nuclear genome

**DOI:** 10.1101/643056

**Authors:** Timothy M. Healy, Ronald S. Burton

## Abstract

Oxidative phosphorylation requires gene products encoded in both the nuclear and mitochondrial genomes, and is the primary source of cellular energy in eukaryotes. As a result, functional integration between the genomes is essential for efficient ATP generation in these organisms. Although within populations this integration is presumably maintained by coevolution, both the importance of coevolution in speciation and mitochondrial disease, and the strength of selection for maintenance of coevolved genotypes are widely questioned. In this study, we crossed populations of the intertidal copepod, *Tigriopus californicus*, to disrupt putatively coevolved mitonuclear genotypes in reciprocal F_2_ hybrids. We utilized inter-individual variation in developmental rate, a proxy for fitness, among these hybrids to assess the strength of selection imposed on the nuclear genome by alternate mitochondrial genotypes. There was substantial variation in developmental rate among hybrid individuals, and *in vitro* ATP synthesis rates of mitochondria isolated from high fitness hybrids were approximately twice those of mitochondria isolated from low fitness individuals. Furthermore, we used Pool-seq to reveal large deviations in nuclear allele frequencies in hybrids, which favored maternal alleles in only high fitness individuals of each reciprocal cross. Therefore, our most fit hybrids had partial recovery of coevolved genotypes, indicating that mitonuclear effects underlie individual-level variation in developmental rate and that inter-genomic compatibility is critical for high fitness. These results demonstrate that mitonuclear interactions have profound impacts on both physiological performance and the evolutionary trajectory of the nuclear genome.

## Introduction

Oxidative phosphorylation in the mitochondria is central to the functioning of essentially all eukaryotic cells, and thus is critical for the majority of complex life (Rand et al., 2004; Lane, 2005; Wallace, 2010a; Hill, 2015). Over evolutionary time most mitochondrial genes have translocated to the nucleus, but a small number that are necessary for ATP generation are still encoded within the mitochondria: typically 13 protein coding, 2 rRNA and 22 tRNA genes in bilaterian animals (Levin et al., 2014). These genes require functional interactions with nuclear-encoded proteins, and thus mitochondrial performance relies upon integration between the nuclear and mitochondrial genomes (Rand et al., 2004; Lane, 2005; Hill, 2015). Consequently, there is predicted to be strong selection for mitonuclear compatibility between interacting genes (i.e., coevolution) in isolated populations and species (Sloan et al., 2017; Hill et al., 2018).

Despite the likelihood of strong selection for compatible mitonuclear genotypes, coevolved mitochondrial and nuclear genes may be disassociated by hybridization when isolated populations experience secondary contact (Burton & Barreto, 2012). Mismatches between mitochondrial-encoded and nuclear-encoded alleles can have profound negative phenotypic consequences across many traits (i.e., hybrid breakdown), ranging from diseases in humans (Wallace, 2010b) to life-history effects in invertebrates (Burton, 1990; Ellison & Burton, 2008b; Meiklejohn et al., 2013). These examples of Bateson-Dobzhansky-Muller incompatibilities (Bateson, 1909; Dobzhansky, 1937; Muller, 1942) may have important implications for post-zygotic isolation between species (Gershoni et al., 2009; Hill, 2016, 2019), and mitochondrial replacement therapies in humans (Reinhardt et al., 2013). However, the ubiquity and relevance of these implications have been questioned for humans specifically (Eyre-Walker, 2017), and for eukaryotes generally (Sloan et al., 2017). Therefore, determining the extent to which mitonuclear interactions influence evolution of the nuclear genome, and the degree to which inter-genomic incompatibilities result in negative fitness consequences are critical for understanding the role of mitochondrial DNA in shaping the physiological performance and evolution of eukaryotes.

In the current study, we address these issues using hybrids between a San Diego, CA (SD) and a Santa Cruz, CA (SC) population of the intertidal copepod *Tigriopus californicus*. This species is found in supralittoral tidepools along the west coast of North America from Baja California, Mexico to Alaska, USA with extremely low gene flow between isolated populations on different rocky outcrops (Burton, 1997). This isolation has led to high levels of genetic divergence among populations (Burton & Lee, 1994; Burton 1997; Edmands, 2001; Peterson et al., 2013; Pereira et al., 2016 Barreto et al., 2018), and F_2_ hybrids from inter-population laboratory crosses typically display breakdown of mitochondrial ATP synthesis capacities and several fitness-related life-history traits, including fecundity and developmental rate (Burton, 1990; Edmands, 1999; Ganz & Burton, 1995; Ellison & Burton, 2008b, Burton et al., 2006). This loss of performance is recovered by backcrossing hybrids to the maternal, but not the paternal, parental population (Ellison & Burton, 2008b), which, since mitochondrial DNA is maternally inherited (Burton & Barreto, 2012), clearly implicates a role for mitonuclear interactions in hybrid breakdown in this species. Here, we reasoned that if there is strong selection for mitonuclear matching throughout ontogeny (Hill et al., 2018), then there should be clear physiological and genetic associations with variation in fitness-related traits among F_2_ hybrids. Thus, we utilized inter-individual differences in developmental rate and ATP synthesis rate in combination with Pool-seq to assess the importance of, and strength of selection for, coevolved mitonuclear genomes in eukaryotes.

## Materials and methods

### Copepod collection and culturing

Adult copepods were collected with fine-mesh dip nets and large plastic pipettes from splashpools near San Diego, CA (SD; 32° 45′ N, 117° 15′ W) and Santa Cruz, CA (SC; 36° 56′ N, 122° 02′ W) in the spring of 2018. Collected animals were transported back to Scripps Institution of Oceanography in 1 L plastic bottles containing water collected from the tidepools. Collections were split into approximately fifteen 200 mL laboratory cultures that were established in 400 mL glass beakers, and held across four incubators for laboratory acclimations. Acclimation conditions (20 °C, 36 ppt and 12:12 light:dark) were maintained for at least one month (approximately one generation) prior to the start of all experiments. Copepods consumed natural algal growth in the cultures as well as a mixture of ground fish flakes and powdered Spirulina that were fed to each culture ad lib.

### Inter-population crosses

Prior to mating, male *T. californicus* clasp juvenile females forming a breeding pair (Burton, 1985). Females mate only once, and thus separation of precopulatory breeding pairs allows the isolation of virgin females for experimental crosses (Burton, 1985). Two sets of reciprocal crosses were made between SD and SC copepods. First, for assessment of variation in ATP synthesis rates (see below), 40 pairs of each population were gently teased apart with a needle (e.g., Burton et al., 1981), and males of one population were combined with females of the other population in 10 cm petri dishes containing ~60 mL of filtered seawater. Copepods were allowed to pair, and the dishes were monitored for the appearance of gravid females, which, when observed, were moved to a new dish. These females were allowed to produce multiple egg sacs each in the new dish, and were removed once F_1_ offspring were visible. F_1_ offspring matured and haphazardly formed breeding pairs. Gravid females were again moved to a new dish, and were monitored until mature (red) F_2_ egg sacs were observed. Second, for isolation of DNA for Pool-seq (see below), 120 pairs of each population were separated, and reciprocal F_2_ hybrid egg sacs were obtained as described above. Throughout the experimental crosses holding conditions and feeding routines were the same as those for the initial laboratory acclimations.

### Inter-individual variation in developmental time

Variation in developmental rate among individuals was assessed for both parental and F_2_ hybrid copepods by measurement of time to metamorphosis (e.g., Harada et al., 2019). *T. californicus* development consists of 6 naupliar stages, 5 copepodid stages and the final adult stage (Tsuboko-Ishii & Burton, 2018). The majority of stages are visually cryptic; however, there is a substantial metamorphosis between the final naupliar stage and initial copepodid stage (i.e., copepodid stage I), which can be observed through a microscope. To score inter-individual differences in developmental rate, gravid females with red egg sacs were pipetted onto filter paper, egg sacs were removed with a fine needle, and dissected egg sacs were placed in filtered seawater in 6-well plates (≤ 4 per well). This procedure synchronizes hatching as dissected mature egg sacs hatch overnight. Offspring were fed Spirulina, and were monitored daily for the appearance of copepodids. Days post hatch (dph) to metamorphosis was scored for all individuals, and copepodids were moved to fresh petri dishes after scoring. In total, offspring from 68 SD egg sacs, 58 SC egg sacs, 352 F_2_ SD♀xSC♂ egg sacs (205 for ATP assays and 147 for Pool-seq) and 314 F_2_ SC♀xSD♂ egg sacs (115 for ATP assays and 199 for Pool-seq) were scored.

### ATP synthesis rates

F_2_ hybrid copepodids were divided into four developmental time groups: 8-10, 11-13, 14-16 and ≥17 dph to metamorphosis. Development was allowed to continue, and adults from the 8-10, 11-13 and ≥17 dph groups were used for assessment of maximal mitochondrial ATP synthesis rates as in Harada et al. (2019). In brief, for each reciprocal cross, 6 pools of 6 adults from each developmental group were moved to petri dishes with fresh filtered seawater and no food overnight. Each pool of copepods was then homogenized in 800 μL of ice-cold homogenization buffer (400 mM sucrose, 100 mM KCl, 70 mM HEPES, 3 mM EDTA, 6 mM EGTA, 1% BSA, pH 7.6) in 1 mL teflon-on-glass homogenizers. Homogenates were transferred to 1.5 mL microcentrifuge tubes, and centrifuged at 1,000 g for 5 min at 4 °C. Supernatants were pipetted to new 1.5 mL tubes, which were then centrifuged at 11,000 g for 10 min at 4 °C. After removal of the supernatants, mitochondrial pellets were resuspended in 55 μL of assay buffer (560 mM sucrose, 100 mM KCl, 70 mM HEPES, 10 mM KH_2_PO_4_, pH 7.6). For the ATP synthesis assays, 5 μL of a complex I substrate cocktail (final assay substrate concentrations: 5 mM pyruvate, 2 mM malate and 1 mM ADP) was added to 25 μL of each sample in 0.2 mL strip tubes. This was done twice for each sample: once for the initial ATP concentration determinations and once for the ATP synthesis reactions. For initial ATP measurements, CellTiter-Glo (Promega, Madison, WI), which is used for ATP quantification and prevents additional ATP synthesis, was immediately added to one tube for each sample after substrate additions. For synthesis reactions, the second tube for each sample was incubated at 20 °C for 10 min prior to the addition of CellTiter-Glo. All samples were incubated with CellTiter-Glo at room temperature in the dark for 10 min prior to reading luminescence with a Fluoroskan Ascent® FL (Thermo Fisher Scientific, Waltham, MA). Sample luminescence was compared to an ATP standard curve, and ATP synthesis rate was calculated by subtracting initial ATP concentrations from final ATP concentrations. Protein content in each sample was measured with NanoOrange Protein Quantification Kits according to the manufacturer’s instructions (Thermo Fisher Scientific, Waltham, MA), and was used for ATP synthesis rate normalization. Variation in synthesis rates among groups was assessed by two-way ANOVA with cross and developmental time as factors followed by Tukey post-hoc tests in R v3.4.0 (The R Foundation, Vienna, Austria).

### Genomic sequencing and allele frequency determination

Two developmental groups of F_2_ hybrid copepodids for each reciprocal cross were allowed to develop to adulthood: those that metamorphosed 8-12 dph (“fast developers”) and those that metamorphosed >22 dph (“slow developers”). For each group, 180 adults (approximately equal numbers of females and males) were pooled for DNA isolation by phenol-chloroform extraction (Sambrook & Russell, 2006). Briefly, copepods were rinsed with deionized water and homogenized by hand in 150 μL of Bender buffer (200 mM sucrose, 100 mM NaCl, 100 mM Tris-HCl pH 9.1, 50 mM EDTA, 0.5% SDS) in 1.5 mL microcentrifuge tubes. An additional 250 μL of Bender buffer was added to each sample followed by 100 μg of Proteinase K (Thermo Fisher Scientific, Waltham, MA). Samples were incubated at 56 °C overnight then cooled to room temperature for ~15 min. 25 μg of RNase A (Thermo Fisher Scientific, Waltham, MA) was added to each sample prior to a 37 °C incubation for 30 min, which was followed by addition of 200 μL of 5M potassium acetate and a 10 min incubation on ice. Samples were then centrifuged at 13,000 g for 10 min at 4 °C, supernatants were transferred to 2.0 mL microcentrifuge tubes, and 400 μL of UltraPure™ Buffer-Saturated Phenol (Thermo Fisher Scientific, Waltham, MA) and 400 μL of OmniPur® Chloroform (EMD Millipore Corporation, Darmstadt, Germany) were added to each supernatant. The phenol-chloroform mixtures were then gently mixed for 1 min, and centrifuged at 20,000 g for 5 min at 4 °C. Aqueous phases were transferred to new 2.0 mL tubes, and organic phases were back-extracted as above to maximize DNA yield. 400 μL of chloroform was again added to each aqueous phase for re-extraction: samples were centrifuged at 15,000 g for 1 min at 4 °C, aqueous phases were transferred to new 2.0 mL tubes, and again organic phases were back-extracted repeating the above procedure. 1,200 μL of ice-cold 95% ethanol was added to each aqueous phase; tubes were incubated at −20 °C for 1 h to facilitate DNA precipitation, and then centrifuged at 16,000 g for 20 min at 4 °C. 95% ethanol was removed by pipette, and all pellets for a sample (from back-extractions etc.) were combined in 1,000 μL of ice-cold 75% ethanol. Samples were then centrifuged at 16,000 g for 5 min at 4 °C. 75% ethanol was removed by pipette followed by an additional 16,000 g centrifugation for 1 min at 4 °C. Any remaining ethanol was removed and samples were dried in air for 20 min then resuspended in UltraPure™ Distilled Water (Thermo Fisher Scientific, Waltham, MA). DNA isolations were quantified with a Qubit® 2.0 Fluorometer and a dsDNA HS assay kit according to the manufacturer’s instructions (Thermo Fisher Scientific, Waltham, MA).

Approximately 1 μg of genomic DNA for each pool was sent to Novogene Co., Ltd. (Sacramento, CA) for whole-genome 150 bp paired-end sequencing on a NovaSeq 6000 (Illumina Inc., San Diego, CA). Between 59,765,048 and 74,464,410 paired reads were obtained for each sample, which were trimmed to remove adapter sequences and base pairs with Phred scores less than 25. After trimming, reads with less than 50 bp remaining were removed. BWA MEM v0.7.12 (Li & Durbin, 2009) was used to align the filtered reads to the SD *T. californicus* reference genome v2.1 (Barreto et al., 2018) and an updated SC reference genome, which was prepared as described in Barreto et al. (2018) and Lima et al. (2019). Prior to read mapping the references were equalized such that any “N” position in one reference was also an “N” in the other reference. Mapping hybrid sample reads to both parental references allows calculation of average allele frequency estimates between the mappings, which accounts for mapping biases between matched and mismatched allelic reads (Lima & Willett, 2018). Read mappings with MAPQ scores less than 20 were discarded, resulting in average genome coverage values between 65X and 83X for all sample-to-reference combinations (Tables 1, 2). 2,768,859 biallelic single nucleotide polymorphisms (SNPs) that are fixed between the SD and SC populations were identified using the previously published methods (Lima & Willett, 2018; Lima et al., 2019) with population-specific sequencing reads obtained from Barreto et al. (2018). Briefly, population-specific reads were mapped to the other population’s reference genome, and variant loci with minor allele frequencies of 0 in both mappings were kept as fixed inter-population SNPs. Sample allele frequencies at these SNPs were determined using PoPoolation2 (Kofler et al., 2011) for all sites that had a minimum coverage of at least 4 for the minor allele in the mappings to both parental reference genomes (as in Lima et al., 2019). Estimated allele frequencies for each sample were averaged between the two mappings to account for mapping biases, and mean allele frequencies were calculated for non-overlapping 250 kb windows along each chromosome, which reduces noise in allele frequency estimates as a single generation of recombination between SD and SC chromosomes (which occurs only in males in *T. californicus* [Burton et al., 1981]) is not expected to break apart large chromosomal blocks in F_2_ hybrids (Lima & Willett, 2018).

**Table 1:** SD♀xSC♂ sequencing and allele frequency summary.

| Chromosome | Number of 250kb windows | Number of SNPs | SNPs per window<sup>1</sup> | KS test *p*-value | Developmental time (dph) | Average coverage | SD allele frequency for 250 kb windows |  |  |
| --- | --- | --- | --- | --- | --- | --- | --- | --- | --- |
| | | | | | | | $\mu$ | $q_{0.1}$ | $q_{0.9}$ |
| One | 66 | 263,305 | 3989 ± 571 | 1.6 x 10<sup>-4</sup> * | 8-12 | 66X | 0.555 | 0.546 | 0.565 |
|  |  |  |  |  | >22 | 77X | 0.470 | 0.432 | 0.493 |
| Two | 61 | 221,662 | 3634 ± 920 | 0.58 | 8-12 | 66X | 0.543 | 0.512 | 0.582 |
|  |  |  |  |  | >22 | 78X | 0.523 | 0.512 | 0.535 |
| Three | 59 | 223,269 | 3784 ± 987 | 1.6 x 10<sup>-4</sup> * | 8-12 | 67X | 0.578 | 0.560 | 0.596 |
|  |  |  |  |  | >22 | 78X | 0.498 | 0.485 | 0.516 |
| Four | 54 | 205,183 | 3800 ± 928 | 5.8 x 10<sup>-4</sup> * | 8-12 | 66X | 0.554 | 0.529 | 0.577 |
|  |  |  |  |  | >22 | 78X | 0.423 | 0.412 | 0.435 |
| Five | 66 | 256,052 | 3880 ± 645 | 1.6 x 10<sup>-4</sup> * | 8-12 | 65X | 0.573 | 0.560 | 0.584 |
|  |  |  |  |  | >22 | 76X | 0.519 | 0.503 | 0.534 |
| Six | 61 | 223,748 | 3668 ± 1086 | 5.8 x 10<sup>-4</sup> * | 8-12 | 63X | 0.473 | 0.460 | 0.487 |
|  |  |  |  |  | >22 | 76X | 0.527 | 0.509 | 0.546 |
| Seven | 66 | 244,443 | 3703 ± 817 | 0.02 | 8-12 | 65X | 0.499 | 0.488 | 0.510 |
|  |  |  |  |  | >22 | 76X | 0.482 | 0.458 | 0.510 |
| Eight | 63 | 236,290 | 3750 ± 717 | 0.58 | 8-12 | 65X | 0.475 | 0.462 | 0.495 |
|  |  |  |  |  | >22 | 77X | 0.481 | 0.460 | 0.511 |
| Nine | 63 | 231,348 | 3672 ± 751 | 0.58 | 8-12 | 65X | 0.481 | 0.462 | 0.500 |
|  |  |  |  |  | >22 | 77X | 0.490 | 0.477 | 0.499 |
| Ten | 65 | 252,232 | 3880 ± 696 | 0.66 | 8-12 | 67X | 0.516 | 0.497 | 0.549 |
|  |  |  |  |  | >22 | 79X | 0.516 | 0.501 | 0.536 |
| Eleven | 63 | 225,229 | 3575 ± 837 | 0.66 | 8-12 | 65X | 0.469 | 0.455 | 0.490 |
|  |  |  |  |  | >22 | 77X | 0.470 | 0.458 | 0.485 |
| Twelve | 72 | 186,098 | 2585 ± 1360 | 6.2 x 10<sup>-3</sup> | 8-12 | 67X | 0.514 | 0.486 | 0.538 |
|  |  |  |  |  | >22 | 78X | 0.486 | 0.462 | 0.507 |

<sup>1</sup> $\mu \pm \sigma$ ; \* significant after Bonferroni correction

**Table 2:** SC♀xSD♂ sequencing and allele frequency summary.

| Chromosome | Number of 250kb windows | Number of SNPs | SNPs per window<sup>1</sup> | KS test *p*-value | Developmental time (dph) | Average coverage | SC allele frequency for 250 kb windows |  |  |
| --- | --- | --- | --- | --- | --- | --- | --- | --- | --- |
| | | | | | | | $\mu$ | $q_{0.1}$ | $q_{0.9}$ |
| One | 66 | 263,305 | 3989 ± 571 | 1.6 x 10<sup>-4</sup> * | 8-12 | 67X | 0.582 | 0.563 | 0.598 |
|  |  |  |  |  | >22 | 80X | 0.492 | 0.481 | 0.504 |
| Two | 61 | 221,662 | 3634 ± 920 | 5.8 x 10<sup>-4</sup> * | 8-12 | 67X | 0.586 | 0.545 | 0.614 |
|  |  |  |  |  | >22 | 80X | 0.470 | 0.448 | 0.493 |
| Three | 59 | 223,269 | 3784 ± 987 | 1.6 x 10<sup>-4</sup> * | 8-12 | 69X | 0.533 | 0.505 | 0.553 |
|  |  |  |  |  | >22 | 81X | 0.456 | 0.446 | 0.472 |
| Four | 54 | 205,183 | 3800 ± 928 | 5.8 x 10<sup>-4</sup> * | 8-12 | 68X | 0.546 | 0.522 | 0.570 |
|  |  |  |  |  | >22 | 80X | 0.471 | 0.454 | 0.489 |
| Five | 66 | 256,052 | 3880 ± 645 | 1.6 x 10<sup>-4</sup> * | 8-12 | 66X | 0.609 | 0.593 | 0.627 |
|  |  |  |  |  | >22 | 78X | 0.492 | 0.481 | 0.505 |
| Six | 61 | 223,748 | 3668 ± 1086 | 0.58 | 8-12 | 67X | 0.486 | 0.468 | 0.505 |
|  |  |  |  |  | >22 | 78X | 0.488 | 0.476 | 0.500 |
| Seven | 66 | 244,443 | 3703 ± 817 | 1.6 x 10<sup>-4</sup> * | 8-12 | 68X | 0.486 | 0.472 | 0.498 |
|  |  |  |  |  | >22 | 80X | 0.451 | 0.435 | 0.464 |
| Eight | 63 | 236,290 | 3750 ± 717 | 5.8 x 10<sup>-4</sup> * | 8-12 | 67X | 0.556 | 0.542 | 0.570 |
|  |  |  |  |  | >22 | 79X | 0.494 | 0.478 | 0.513 |
| Nine | 63 | 231,348 | 3672 ± 751 | 8.2 x 10<sup>-3</sup> | 8-12 | 68X | 0.480 | 0.468 | 0.493 |
|  |  |  |  |  | >22 | 80X | 0.503 | 0.487 | 0.518 |
| Ten | 65 | 252,232 | 3880 ± 696 | 0.09 | 8-12 | 69X | 0.505 | 0.488 | 0.520 |
|  |  |  |  |  | >22 | 81X | 0.516 | 0.496 | 0.532 |
| Eleven | 63 | 225,229 | 3575 ± 837 | 1.6 x 10<sup>-4</sup> * | 8-12 | 69X | 0.493 | 0.480 | 0.508 |
|  |  |  |  |  | >22 | 79X | 0.547 | 0.532 | 0.561 |
| Twelve | 72 | 186,098 | 2585 ± 1360 | 4.1 x 10<sup>-5</sup> * | 8-12 | 71X | 0.475 | 0.465 | 0.498 |
|  |  |  |  |  | >22 | 83X | 0.444 | 0.418 | 0.470 |

<sup>1</sup> $\mu \pm \sigma$ ; \* significant after Bonferroni correction

Deviations in F_2_ allele frequencies associated with mitonuclear effects between fast and slow developers were detected for each cross as in Lima et al. (2019). First, average maternal allele frequencies were calculated for non-overlapping 2 Mb blocks along each chromosome (7-9 blocks per chromosome), and Kolmogorov-Smirnov (KS) tests in R v3.4.0 were used to assess if the allele frequencies across each chromosome in each pool were drawn from the same distribution. False-discovery rate associated with multiple tests was accounted for by Bonferroni correction of α = 0.05. Second, for chromosomes with significant KS test results, the tenth (*q*_0.1_) and ninetieth (*q*_0.9_) quantiles of the allele frequencies calculated over 250 kb blocks were compared between fast and slow developers. Lack of overlap between the quantiles is consistent with a potential mitonuclear effect on developmental rate. These potential effects were then resolved by comparisons between the reciprocal crosses; higher maternal allele frequencies in the fast developers than the slow developers in both reciprocals or in one reciprocal with no allele deviations in the other reciprocal are consistent with effects of mitonuclear matching. In contrast, higher paternal alleles frequencies in fast developers using the same comparisons are consistent with mitonuclear mismatching. Third, allele frequency deviations greater than or equal to ±0.05 were identified, as this minimum deviation has been suggested as a threshold for chromosomal regions most likely to contain genes involved in mitonuclear effects using these methods (Lima & Willett, 2018). These analyses were repeated comparing SD allele frequencies between the reciprocal crosses for the fast and slow pools separately as an alternative test of allele deviations consistent with mitonuclear incompatibilities. In fast developers, greater frequencies of the SD allele in the SD♀xSC♂ than the SC♀xSD♂ cross are consistent with effects of mitonuclear matching. In contrast, expected patterns in slow developers are more challenging to predict. For example, lower frequencies of the SD allele in the slow developing SD♀xSC♂ copepods than in the slow developing SC♀xSD♂ copepods would be consistent with effects of mitonuclear incompatibilities. However, because different sites of mismatches could independently cause similar negative phenotypic effects and individuals with many mismatches might be expected to fail development at early stages, it is also possible that few allelic deviations would be expected in slow developers even if mitonuclear interactions affected developmental rate. Finally, effects of nuclear genetic variation alone were assessed by determining the chromosomes for which allele frequencies deviated in the same direction from 0.5 in both fast developing reciprocal pools or both slow developing reciprocal pools, such that *q*_0.1_ and *q*_0.9_ did not overlap with 0.52 or 0.48 in either reciprocal. These boundaries are likely conservative quantile limits for neutral variation in allele frequency estimates using these methods (Lima & Willett, 2018; Lima et al., 2019).

## Results

Developmental rates were similar in both parental populations of *T. californicus* with metamorphosis occurring approximately 8-22 days post hatch (dph) for ~98% of nauplii (maximum dph of 29 and 24 for SD and SC, respectively; Fig. 1A). In contrast, the distributions of developmental time among F_2_ hybrids from both reciprocal crosses demonstrated a substantial skew favoring higher dph to metamorphosis compared to the parental populations, which is consistent with hybrid breakdown (Fig. 1B). In both crosses, metamorphosis was observed 8-30 dph with 8 out of 473 SD♀xSC♂ nauplii and 245 out of 1,242 SC♀xSD♂ nauplii still present on day 30, which were scored as >30 dph. Preliminary data suggested that the majority of nauplii underwent metamorphosis 9-16 dph, and thus F_2_ hybrids were split into 8-10, 11-13 and ≥17 dph groups to assess maximal ATP synthesis rates. Complex I-fueled ATP synthesis rates were significantly affected by both cross (F_1,30_ = 11.32; *P* = 2.1 x 10^−3^) and developmental group (F_2,30_ = 13.44; *P* = 6.8 x 10^−5^) with no interaction between factors (F_2,30_ = 0.44; *P* = 0.65), and post-hoc tests indicated that 8-10 dph copepods had higher ATP synthesis rates than ≥17 dph copepods in both crosses (*P* ≤ 0.04; Fig. 1C). In SC♀xSD♂ hybrids, 11-13 dph copepods had ATP synthesis rates that were similar to those of 8-10 dph copepods (*P* = 0.52) and higher than those of ≥17 dph copepods (*P* = 0.01). In contrast, 11-13 dph SD♀xSC♂ copepods had intermediate synthesis rates compared to 8-10 and ≥17 dph copepods (*P* ≥ 0.13 for both comparisons; Fig. 1C).

**Fig. 1.**
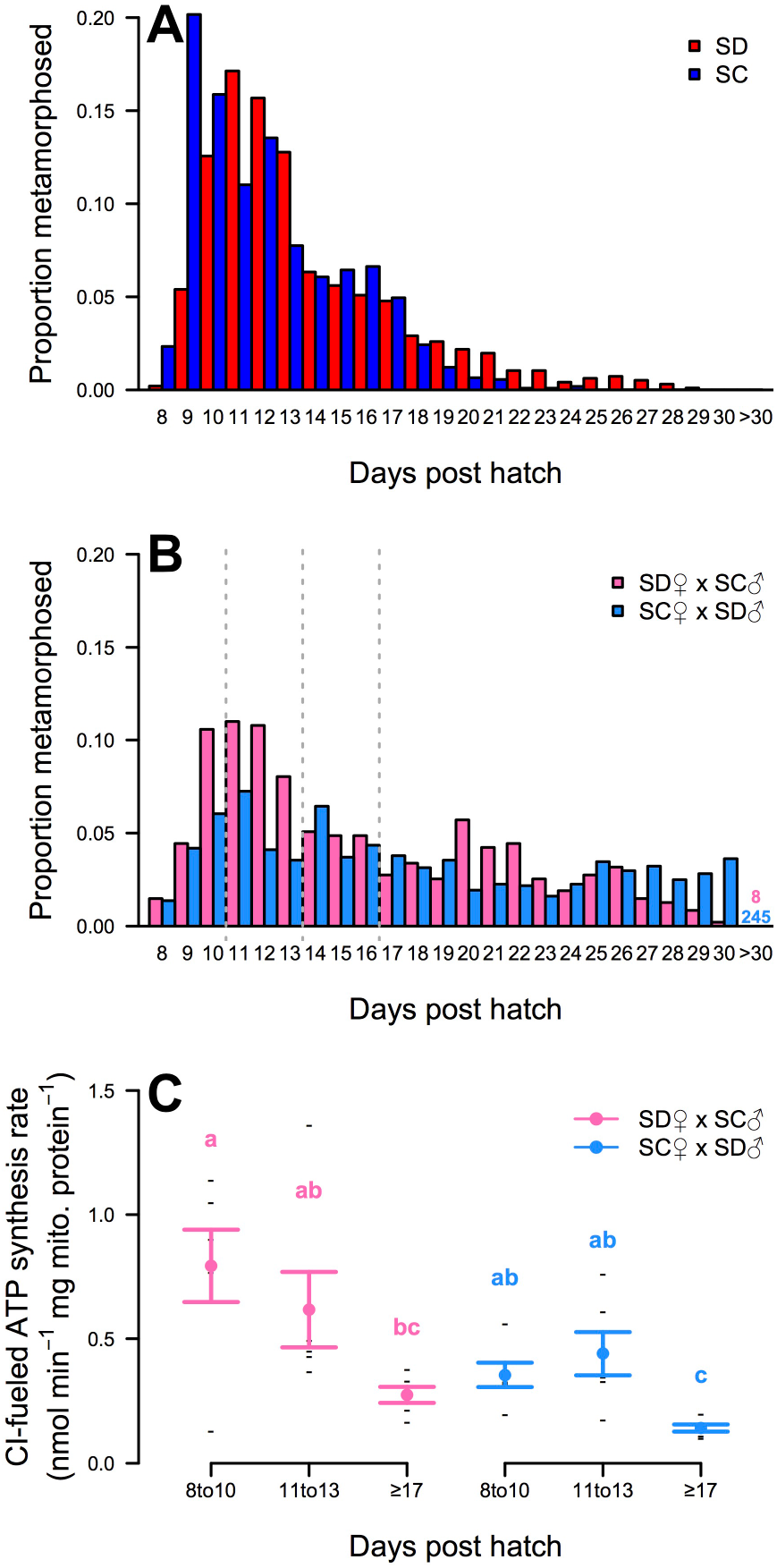
Developmental time to metamorphosis for *T. californicus* nauplii as proportion of individuals; A: SD (red; *N* = 963) and SC (blue; *N* = 1,071), and B: SD♀xSC♂ (pink; *N* = 473) and SC♀xSD♂ (light blue; *N* = 1,242) F_2_ hybrids. F_2_ hybrids were split by developmental time (dashed grey lines) and maximal complex I (CI)-fueled ATP synthesis rates were measured for adults that metamorphosed 8-10, 11-13 and ≥17 dph for both reciprocal crosses (C: mean ± s.e.m. for SD♀xSC♂ - pink and SC♀xSD♂ - light blue; all measurements - black dashes; *N* = 6 per group). Shared lower case letters indicate groups that do not differ significantly.

F_2_ hybrids from a second set of reciprocal crosses were divided into those that metamorphosed 8-12 or >22 dph (fast or slow developers, respectively) to assess parental allele frequency deviations associated with variation in developmental rate. When considering all 2,768,859 SNPs that were fixed between the SD and SC populations, there were shifts towards higher maternal allele frequencies in fast developers, and higher paternal allele frequencies in slow developers in each reciprocal cross (Fig. 2). Biases towards maternal alleles in fast developers became particularly evident when allele frequencies were examined across chromosomes (Fig. 3), as in both reciprocal crosses there were significant frequency shifts favoring coevolved alleles in fast developers across large regions of chromosomes 1, 3, 4 and 5 (*P* ≤ 5.8 x 10^−4^; Figure 3A). Although deviations within a single reciprocal could be consistent with nuclear-only effects, the observation of maternal biases in both crosses clearly suggests involvement of mitonuclear interactions. Patterns consistent with mitonuclear effects were also detected for chromosomes 2 and 8 in SC♀xSD♂ hybrids, as there were biases for SC alleles in fast developers in this cross (*P* = 5.8 x 10^−4^ for both; Fig. 3B), and these alleles were not favored in fast developing SD♀xSC♂ hybrids. Additionally, there were significant potential mitonuclear effects on chromosomes 7 and 12 in the SC♀xSD♂ cross (*P* ≤ 1.6 x 10^−4^ for both; Fig. 3B); however, these patterns were subtle relative to those on other chromosomes. On chromosome 7, SC alleles were more common in fast than in slow developers, but SC allele frequencies did not exceed 0.5 in either pool, and on chromosome 12, trends in allele frequencies suggested a similar pattern to the one observed on chromosome 7, but quantile overlap comparisons did not conclusively resolve the direction of this potential effect. In general, comparisons between the reciprocal crosses for fast or slow developers demonstrated similar mitonuclear effects to those describe above, and there were no clear nuclear-only effects for any chromosome (Fig. 4). Furthermore, these comparisons detected biases for paternal (i.e., mismatched) nuclear alleles in slow developers (for chromosomes 1, 3, 4, 7 and 12; Fig. 4). Despite the overall association between high fitness and coevolved mitonuclear genotypes in our study, allele frequencies across one chromosome in each reciprocal were consistent with the opposite effect as excess paternal alleles were observed in fast developers (chromosome 6 for SD♀xSC♂ and chromosome 11 for SC♀xSD♂; Fig. 3). Yet, taken together these results suggest that mitonuclear interactions are the major genetic factors contributing to inter-individual variation in developmental rate among F_2_ hybrids, and that in the majority of cases at least partial maintenance of coevolved mitonuclear genotypes is critical for performance in this fitness-related trait.

**Fig. 2.**
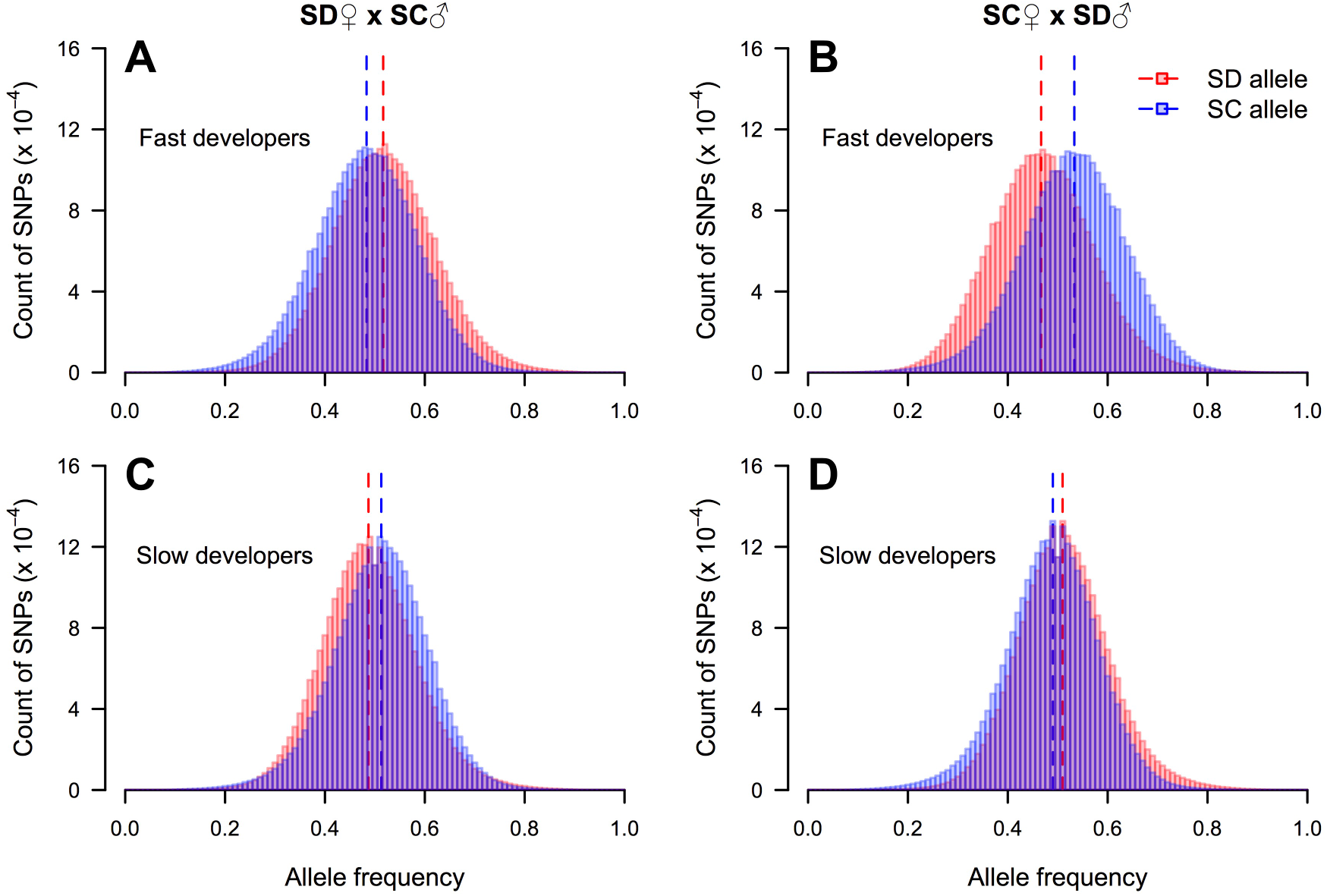
Parental allele frequency histograms (SD - red and SC - blue) for all nuclear single nucleotide polymorphisms (SNPs; *N* = 2,768,859) in fast developing (A: SD♀xSC♂; B: SC♀xSD♂) and slow developing (C: SD♀xSC♂; D: SC♀xSD♂) reciprocal F_2_ hybrids.

**Fig. 3.**
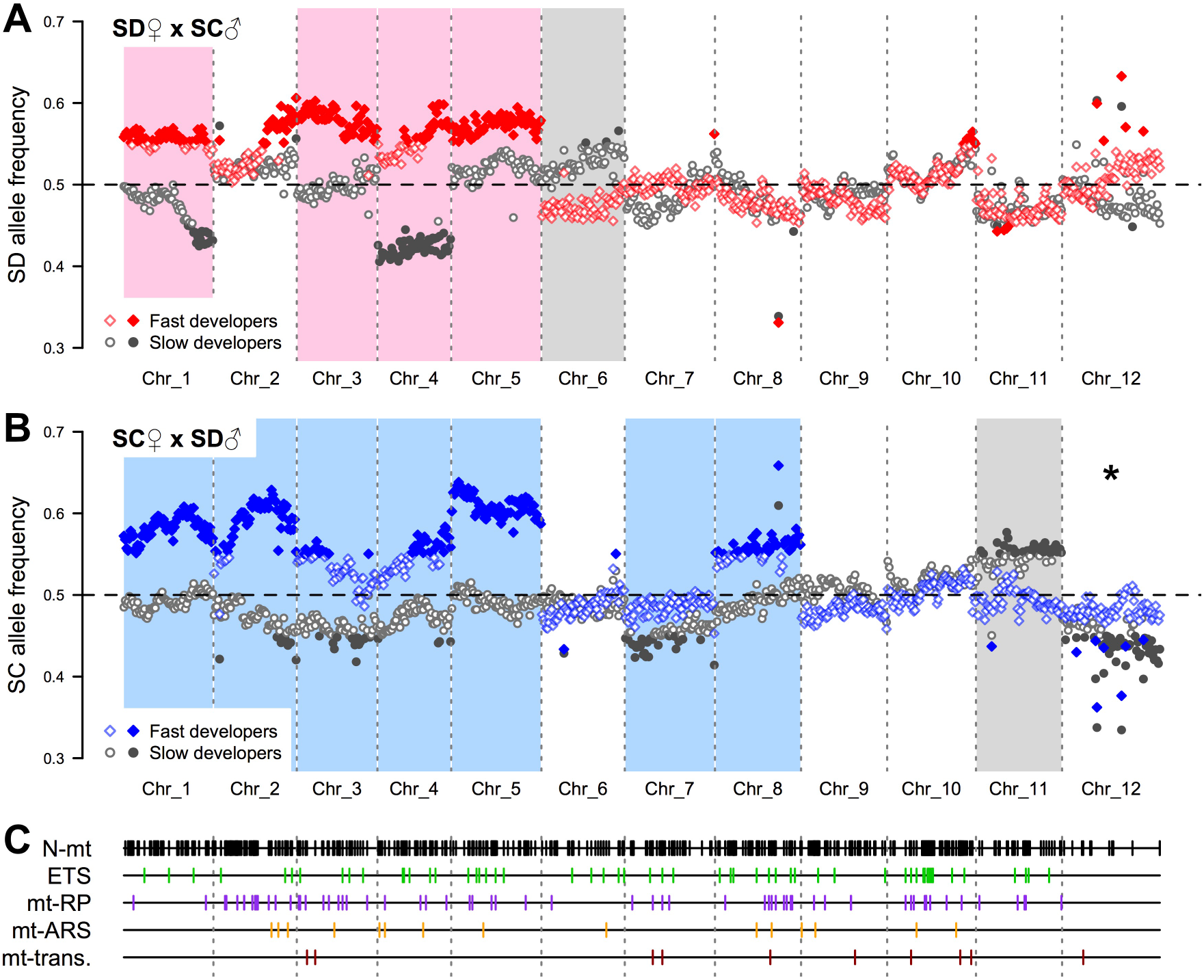
Maternal allele frequencies in SD♀xSC♂ (A) and SC♀xSD♂ (B) fast (red or blue diamonds) and slow (grey circles) developing F_2_ hybrids. Shaded boxes indicate chromosomes with significant deviations consistent with mitonuclear interactions (pink and light blue - maternal bias in fast compared to slow developers; grey - maternal bias in slow compared to fast developers). Filled symbols show allele frequency deviations ≥0.05 from the expected value of 0.5. The asterisk indicates a significant difference in allele frequencies between fast and slow developers that was unresolved by quantile comparisons. Locations of 599 nuclear-encoded mitochondrial genes are displayed in Panel C (all N-mt genes - black; classes of N_O_-mt genes: electron transport system [ETS] - green; ribosomal proteins [mt-RP] - purple; aminoacyl tRNA synthetases [mt-ARS] - orange; transcription and DNA replication [mt-trans.] - dark red).

**Fig. 4.**
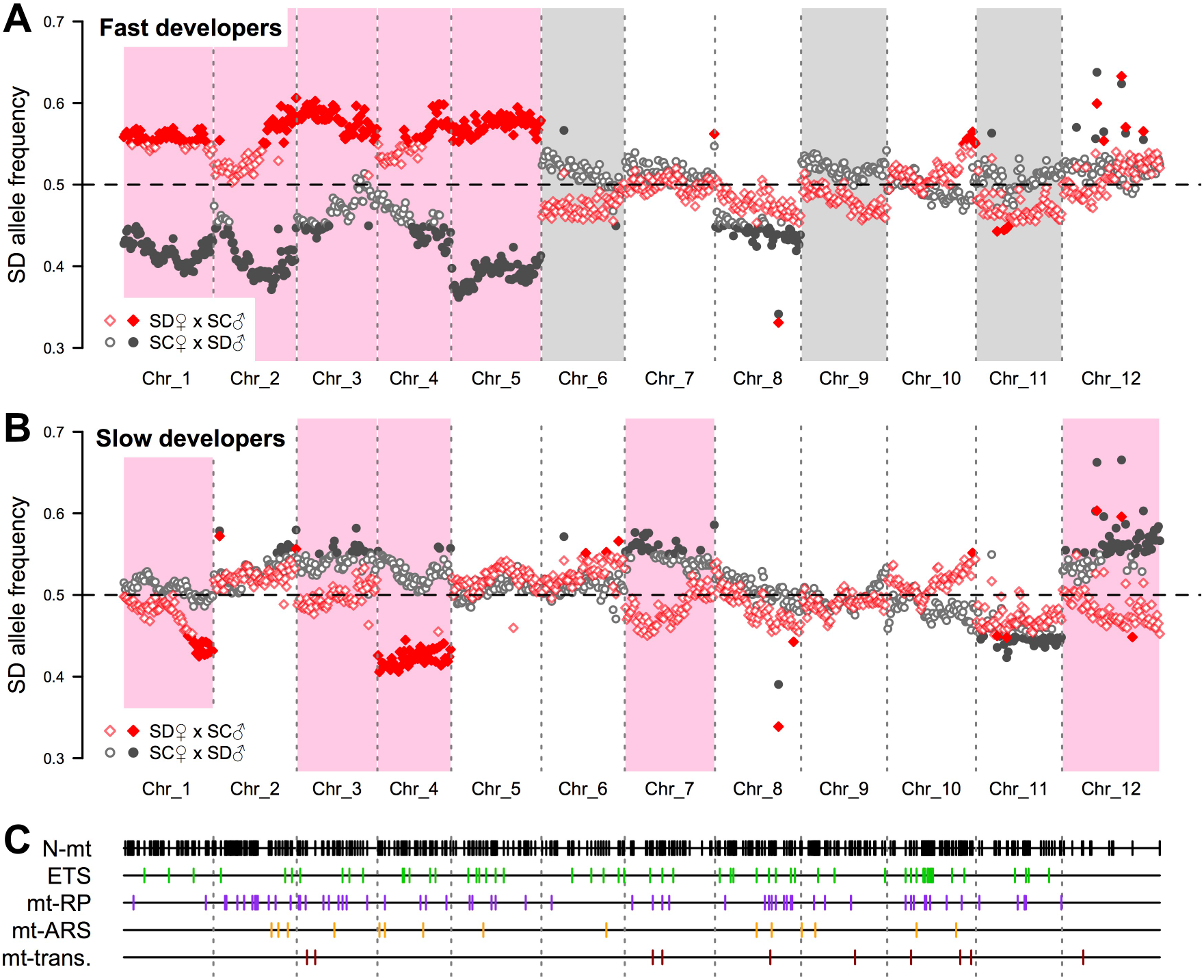
SD allele frequencies in fast (A) and slow (B) developing SD♀xSC♂ (red diamonds) and SC♀xSD♂ (grey circles) F_2_ hybrids. Pink and grey shaded boxes indicate chromosomes with significant deviations consistent with positive effects of mitonuclear matching or mismatching, respectively, on developmental rate. Filled symbols show allele frequency deviations ≥0.05 from the expected value of 0.5. Locations of 599 nuclear-encoded mitochondrial genes are display in Panel C (all N-mt genes - black; classes of N_O_-mt genes: electron transport system [ETS] - green; ribosomal proteins [mt-RP] - purple; aminoacyl tRNA synthetases [mt-ARS] - orange; transcription and DNA replication [mt-trans.] - dark red).

## Discussion

Mitochondrial DNA contains relatively few genes, but because of the functional products encoded by these genes and their interactions with nuclear gene products, differences in mitochondrial genotype are predicted to exert strong selection pressures on the nuclear genome throughout ontogeny (Hill et al., 2018). In the current study, we demonstrate substantial consequences of mitonuclear interactions on developmental rate, ATP synthesis rate, and nuclear allele frequencies in hybrids that are consistent with strong selection for compatible interactions within even a single generation. In our reciprocal crosses, coevolved nuclear alleles that matched alternate mitochondrial genotypes were favored on at least four of the twelve chromosomes in high fitness F_2_ hybrids. Relative to previous studies in *T. californicus* hybrids (Pritchard et al., 2011; Foley et al., 2013; Lima et al., 2019), this clear pattern towards partial recovery of coevolved mitonuclear genotypes is most likely a consequence of selecting individuals based on variation in a fitness-related trait that has been correlated with mitochondrial performance in this species (Ellison & Burton, 2006). Average chromosome-wide allele frequency deviations favoring coevolved alleles ranged from 0.033 to 0.109 with some regions favoring these alleles by ~0.138 (Figs. 3, 4). F_1_ hybrids between SD and SC (heterozygous across all fixed SNPs) generally show enhanced fitness compared to parentals (Ellison & Burton, 2008b), and there is little evidence for selection against heterozygous F_2_ hybrids (Pritchard et al., 2011; Foley et al., 2013). Therefore, it is likely that the major allele frequency deviations in the current study are consequences of negative effects associated with one of the two possible homozygous genotypes. As a result, given Mendelian segregation ratios of 1:2:1, the most extreme biases for maternal alleles in our study are likely indicative of approximately 67-87% deficits of homozygous paternal genotypes in fast developing F_2_ hybrids on some regions of these chromosomes. Thus, our data demonstrate strong selection favoring mitonuclear compatibility. Additionally, despite previous detection of nuclear-only effects on intrinsic selection for survival in *T. californicus* hybrids (Lima et al., 2019), we observed no clear allele frequency deviations consistent with nuclear-only effects on developmental rate, which also suggests that mitonuclear incompatibilities are key genetic mechanisms resulting in loss of fitness in these hybrids.

Previous studies have demonstrated at least three candidate mechanisms involved in coevolution in *T. californicus*: electron transport system complex activities (Willett & Burton, 2001, 2003; Rawson & Burton, 2002; Harrison & Burton, 2005; Ellison & Burton, 2008b), mitochondrial transcription (Ellison & Burton, 2008a) and mitonuclear ribosomal interactions (Barreto & Burton, 2012). Yet, these studies do not directly reveal the number or relative importance of mitonuclear incompatibilities contributing to hybrid breakdown in this species. In comparison, our Pool-seq approach resolves essentially the full genomic architecture of breakdown of developmental rate in hybrids between SD and SC. The substantial biases for maternal alleles across multiple genomic regions in our most fit hybrids clearly indicate a polygenic basis for mitonuclear coevolution, which may be attributable to the high level of divergence between the mitochondrial genomes of these populations (21.7%; Barreto et al., 2018). However, because of the mitochondrion’s central role in metabolism, even minor disruption of mitonuclear interactions can have major fitness effects (Burton & Barreto, 2012, Hill et al., 2018). For example, mutations in a single nuclear-encoded mitochondrial tRNA synthetase and one mitochondrial tRNA lead to mitochondrial dysfunction in *Drosophila* hybrids (Meiklejohn et al., 2013). The allele frequency variation in our slow developing hybrids is likely consistent with large effects of few interactions in *T. californicus* as well. Despite an approximately two-fold reduction in both developmental rate and ATP synthesis rate (Fig. 1), strong deviations favoring paternal alleles in slow developers were largely absent in our study (with the exception of chromosome 4 in SD♀xSC♂; Figs. 3, 4). Consequently, our data suggest that for any given individual, relatively few incompatible interactions are necessary to result in substantial negative fitness effects.

Of the 1,000-1,500 nuclear-encoded mitochondrial (N-mt) genes in metazoans, at least 180 are expected to have intimate functional interactions with either mitochondrial DNA or mitochondrial-encoded gene products (N_O_-mt genes) (Burton et al., 2013). Barreto et al. (2018) identified 599 putative N-mt genes, including 139 N_O_-mt genes, in the *T. californicus* genome (Figs. 3C, 4C). Although our data begin to resolve which of these candidates may play the largest roles in inter-genomic coevolution, there was little resolution of allele frequency deviations beyond the level of chromosomes in our hybrids. This is likely a consequence of only a single opportunity for inter-population recombination in F_2_ hybrids (Lime & Willett, 2018), or involvement of multiple loci within the same chromosome (Willett et al., 2016). Although N-mt genes were not more common on the chromosomes with biases for coevolved alleles than on other chromosomes in our study, chromosomes demonstrating allelic biases tended to have relatively higher ratios of N_O_-mt genes to other N-mt genes (Fig. 5), which is consistent with a disproportionate role for N_O_-mt genes in mitonuclear interactions.

**Fig. 5.**
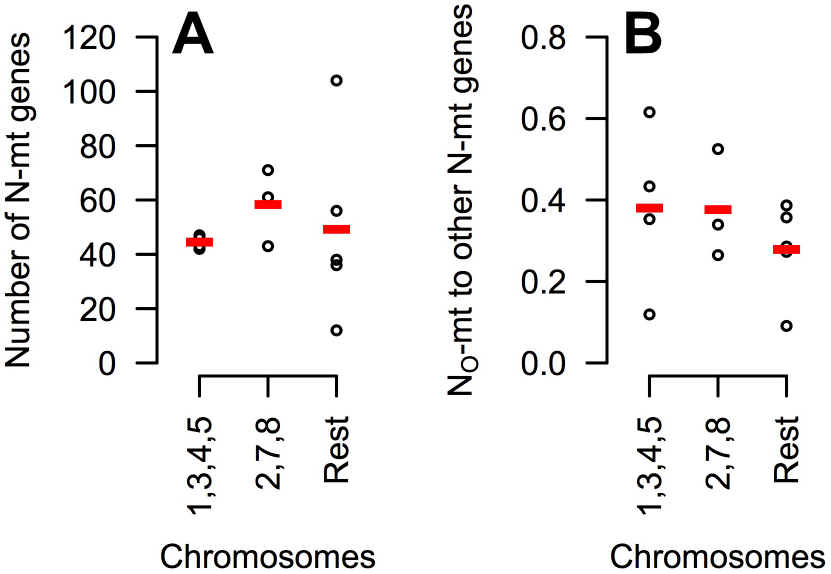
Number of nuclear-encoded mitochondrial (N-mt) genes (A) and ratios of putative interacting (N_O_-mt genes) to other N-mt genes (B) for all twelve *T. californicus* chromosomes. Chromosomes are grouped by those that were consistent with effects of mitonuclear matching on development detected and resolved in both the SD♀xSC♂ and SC♀xSD♂ crosses (1, 3, 4 and 5), in only the SC♀xSD♂ cross (2, 7 and 8), or in neither of the crosses (6, 9, 10, 11 and 12): individual chromosome values - black circles; mean values - red dashes.

Taken together, our data demonstrate strong selection against disassociation of coevolved genes following hybridization, and conversely, strong selection for inter-genomic compatibility within populations and species. These effects of mitonuclear interactions are sufficiently strong in *T. californicus* that selection for rapid development within a single generation identified key sites of mitonuclear interactions across the genome. Mitonuclear coevolution in this species may be exceptionally strong (Burton et al., 2006), but our results also suggest that even small numbers of mitonuclear incompatibilities may result in large losses of fitness. Thus, the findings of the current study are consistent with suggestions that inter-genomic incompatibilities may play a significant role in establishing reproductive isolation between populations (Gershoni et al., 2009; Hill, 2016, 2019). Finally, the potential strength of mitonuclear incompatibilities, their polygenic nature, their range of impacts and their penetrance supports calls for continued caution regarding human mitochondrial replacement therapy.

## Acknowledgments

The current study was funded by a National Science Foundation grant to RSB (DEB1551466). The authors thank Drs. Felipe Barreto and Thiago Lima for advice regarding the sequencing methods and analyses, and Laura Furtado, Antonia Bock and Rebecca Pak for assistance with copepod culturing.

## Author contributions

T.M.H. and R.S.B. designed the experiments in the current study. T.M.H. performed the experiments and analyses, and R.S.B. conceived and supervised the study. T.M.H. prepared the figures, and T.M.H. and R.S.B. wrote the manuscript.

